# Role of *Drosophila* Rap/Fzr (DCdh1) in Retinal Axon Targeting and its Interactions with Loco and Liprin-alpha

**DOI:** 10.1101/2020.05.06.080986

**Authors:** Marta Grońska-Pęski, Tadmiri R. Venkatesh

## Abstract

The development of the wild type *Drosophila* compound eye involves stereotypical targeting of photoreceptor axons to the specific layers of the optic ganglion, medulla and lamina, in the third instar larvae. To test the hypothesis that ubiquitin ligases play an important role during retinal axon targeting we have examined the patterns of axon targeting in the developing eye of the *retina aberrant in pattern (rap/fzr)* mutants. Rap/Fzr is a homolog of mammalian Cdh1, an activator of anaphase promoting complex (APC), a multi-subunit E3 ubiquitin ligase, regulating the cell cycle progression. Previous work has shown that Rap/Fzr is required during eye development for proper cell cycle regulation, glia differentiation and pattern formation. It was also necessary for proper neuromuscular junction development and circadian rhythms. Our results show that Rap/Fzr is required for proper retinal axon targeting in the developing eye. Using *ro-tau-lacZ*, we show that the R2-R5 axons fail to stop in the lamina and mis-target to the medulla levels. Also, mosaic analyses experiments using FLP-FRT and GAL4-UAS techniques show that Rap/Fzr functions in a cell autonomous manner. To test for possible role of other signalling molecules and interactions with Rap/Fzr, we have examined *rap/fzr* axon projection phenotypes in double mutant combinations with the RGS protein, *locomotion defective* (*loco*) mutants and a scaffolding protein, Liprin-α. Our studies suggest that Rap/Fzr is required for proper axon targeting during *Drosophila* visual system development, and the phenotype is enhanced in double mutants with either *loco* or Liprin-α. These results are consistent with other mammalian studies reporting a role of Cdh1 in axon growth and targeting and provides further insights into neuronal functions of the ubiquitin ligase APC/C^Cdh1^.

**Highlights:**

- Loss of *rap/fzr* in the third instar Drosophila eye disc leads to photoreceptor axon overgrowth
- Overexpression of *rap/fzr* leads to photoreceptor axon leads to axon shortening and clumping
- Loss of *Loco^P452^* leads to photoreceptor overgrowth
- Double mutants of *rap* and *loco* or *rap* and *Liprin-α* show axon enhancement of the axon targeting defects in the *Drosophila* third instar larvae eye imaginal discs.

## Introduction

The *Drosophila* eye development is a tractable experimental system for in vivo studies aimed at understanding the mechanisms of cellular pattern formation and their functional abnormalities. The *Drosophila* compound eye is composed of 750 identical subunits called ommatidia. Each ommatidium contains a cluster of eight photoreceptor cells (R cells), six outer R cells (R1-R6) and two inner R cells (R7 and R8) (Ting and Lee, 2007). The photoreceptor axons project in a stereotypical manner to their target ganglion layers in the optic lobe (Clandinin and Zipursky, 2002). R1-R6 photoreceptor axons target the lamina, the first optic ganglion, and R7 and R8 axons target deeper layers in the optic lobe, the M6 and M3 medulla layers, respectively (Ting and Lee, 2007; Morey et al., 2008). R cell patterning seems to be independent of the visual input and neuronal activity (Clandinin and Zipursky, 2000). The availability of genetic tools to manipulate the developing *Drosophila* eye allows for in vivo studies of biochemical pathways responsible for the stereotypical photoreceptor neurons pattern formations in the brain.

There are several *Drosophila* mutants identified displaying various defects in the targeting of the photoreceptor axons to the optic ganglion, including mistargeting, hypo- and hyper-innervation, and clumping of axons (Martin et al., 1995; Lee et al., 2001; Kaminker et al., 2002; Fan et al., 2005). Other aspects of axonal development such as growth cone morphology, axon growth, or cell-cell interactions, were also affected in many mutants (Berger et al., 2008). To further explore axon targeting mechanisms, we screened *rap, retina aberrant in pattern*, previously reported to show a rough eye external phenotype (Karpilow et al., 1989) and involvement in R1, R6 and R7 photoreceptor cell fate determination (Karpilow et al., 1996), for axon projection defects.

*Rap/Fzr* is a *Drosophila* homolog of the mammalian Cdh1 (D-^*Cdh1*^), a regulatory subunit of APC/C, Anaphase Promoting Complex/Cyclosome. APC/C is a E3 ubiquitin ligase complex that targets substrate proteins for ubiquitination and proteasomal degradation (Yang et al., 2010). Previous studies have shown that D-^*Cdh1*^ (Rap/Fzr) is required for the regulation of mitotic events and pattern formation during compound eye development (Karpilow et al., 1996). *rap/fzr* plays a role in synaptic plasticity and bouton morphology and structure (Wise et al., 2013). A genetic modifier screen identified 39 D-^*Cdh1*^ interacting genes, involved in ubiquitin mediated proteolysis, signal transduction, and transcriptional regulation, one of which was *locomotion defects* (l*oco*) (Kaplow et al., 2007).

*loco* encodes a regulator of G-protein signalling (RGS), an important family of proteins critical in the regulation of signalling mediated by G-coupled receptors (GPCR) (Yu et al., 2005). Loco physically interacts with Rap/Fzr through a recognition binding motif D-box, a destruction binding motif creating a post-translational mechanism of Loco degradation by 26S proteasome (Kaplow et al., 2007; Kaplow et al., 2008).

Loco contains a RGS domain and Go Loco motif, which in *Drosophila* functions as a guanine nucleotide exchange factor (GEF) and is expressed primarily in the neuroblasts and in the lateral and midline glia cells in the central nervous system. Its disruption causes several developmental defects such as improper ensheathing of neuronal axons and blood-brain barrier disruption (Granderath et al., 1999; Granderath and Klambt, 2004). Loco is involved in the formation, extension, and migration of glia cells and the asymmetric cell division of neuroblasts (Yu et al., 2005). The data suggests that Loco is required for glia differentiation and triggers glia migration and differentiation in the brain (Silies et al., 2010). Very little is known about Loco expression in non-glial cells, but it was shown to be involved in the dorso-vental patterning and circadian regulation of gene expression (Claridge-Chang et al., 2001; Pathirana et al., 2001). Whether *loco* plays additional functions in non-neuronal cells such as photoreceptor axon targeting is unknown.

Liprin-α is another potential regulator of axon targeting, previously shown to be required for proper R1-8 targeting (Kaufmann et al., 2002; Choe et al., 2006a; Hofmeyer et al., 2006). Liprin-α is a scaffolding protein capable of mediating biochemical indirect interaction or through adaptor proteins, and is known to interact with APC/C to regulate bouton numbers and other neuronal functions (Teng and Tang, 2005), but their role in axonal targeting is not well understood. Thus, we wanted to investigate whether Liprin-α and *rap* act on the same process of axon targeting to enhance defects in photoreceptor axon targeting.

In the present study, we tested the roles of *rap/fzr* in the third instar *Drosophila* eye-brain complex. We examined the morphology of R1-R8 photoreceptor axons of the *rap* mutants in the optic lobe of the third instar larvae and its interaction with *loco* and Liprin-α. Using UAS/GAL4 system (Duffy, 2002) and FLP/FRT (Xu and Rubin, 1993; Theodosiou and Xu, 1998), overexpression and loss-of-function clones, we show R cell axonal abnormalities in the *rap* mutants such as axonal overgrowth, clumping of R cells and overall disorganization of the axonal projections. Quantification of the *rap* mutants showed allele specific changes in the R cell lengths indicating a possible role of *rap* in the control of axonal extension. Double mutants of *rap* with *liprin-α* or *loco* showed enhancement in the axonal defects suggesting their involvement in proper R cell axonal development and targeting.

## Materials and Methods

### Fly stocks and culture

The stocks were maintained at 25°C on standard corn meal-agar medium with 12 h:12 h light:dark cycles.

### Drosophila Stocks

*rap* mutants alleles were generated by (Pimentel and Venkatesh, 2005) using EMS and X-ray mutagenesis: *rap^x-3^; rap^x-2^; w,rap^3^; rap^E2^; rap^E4^; rap^E6^*. For R2-R6 axonal pattern we used *ro-tau-lacZ* (gift from Iris Salecker).

| **Rap allele** | **Phenotype** | **Classification** | **Mutagen** |
| --- | --- | --- | --- |
| *rap*<sup>x-3</sup> | Rough eye in males | Partial loss of function | x-rays |
| *rap*<sup>x-2</sup> | Rough eye in males | Partial loss of function | x-rays |
| *w,rap*<sup>3</sup> | Rough eye in males | Partial loss of function | p-element |
| *rap*<sup>E2</sup> | Rough eye in males | Partial loss of function | EMS |
| *rap*<sup>E4</sup> | Rough eye in males | Partial loss of function | EMS |
| *rap*<sup>E6</sup> | Rough eye in males | Partial loss of function | EMS |

Overexpression studies were performed with the following stocks: *Repo-Gal4/Tb*, (733) *UAS-fzr II.I*.

For mosaic analysis we used *FRT19, Ubi-GFP, eyFLP1* (gift from Jessica Treisman), (111943) *w^67c23^P{lacW}rap^G0418^P{neoFRT}19A/FM7c; P{ey-FLP.DN}5* (Kyoto Institute of Technology), *hsp.FLP; Act>CD2>Gal4 UAS-GFP/TM6B*.

Balancing stocks used were: *w; If/CyO; MKRS/Tb; w; Tm3,Sb/Tm6,Tb; yw; lwr^3-4(2^)/CyO-Act-GPF* (gift from Shuba Govind).

Bloomington Drosophila Stock Center Stocks Loco-stocks used: *Loco^p452^/Tb*; (10009), *P{ry[+*]=lacZ-un1}loco[rC56]*; (14421), *y[1]; ry[506] P{y[+mDint2] w[BR.E.BR]=SUPor-P}loco[KG02176]/TM3, Sb[1] Ser[1]; (19915), y[1] w[67c23]; P{w[+mC] y[+mDint2]=EPgy2}loco[EY04589]*; (25055), *w[1118]; Df(3R)BSC527/TM6C, Sb[1]*; (26883), *w[*]; P{w[+mC]=EP}loco[GE24954]*.

### Genetic Crosses used

#### A. *rap* alleles and ro-tau lacZ

*rap^E2^/rap^E2^; +/+; +/+ x w/Y; Bl/CyO; ro-tau lacZ/ro-tau lacZ*

*rap ^E4^/rap^E4^; +/+; +/+ x w/Y; Bl/CyO; ro-tau lacZ/ro-tau lacZ*

*rap ^E6^/rap^E6^; +/+; +/+ x w/Y; Bl/CyO; ro-tau lacZ/ro-tau lacZ*

*rap ^x-2^/rap^x-2^; +/+; +/+ x w/Y; Bl/CyO; ro-tau lacZ/ro-tau lacZ*

*rap ^x-3^/rap^x-3^; +/+; +/+ x w/Y; Bl/CyO; ro-tau lacZ/ro-tau lacZ*

*w,rap^3^/w,rap^3^; +/+; +/+ x w/Y; Bl/CyO; ro-tau lacZ/ro-tau lacZ*

*CS/CS; +/+; +/+ x w/Y; Bl/CyO; ro-tau lacZ/ro-tau lacZ*

*w^1118^ /w^1118^; +/+; +/+ x w/Y; Bl/CyO; ro-tau lacZ/ro-tau lacZ*

*+/+; +/+; Loco^P452^/Tbx w/Y; Bl/CyO; ro-tau lacZ/ro-tau lacZ*

#### B. Gain of function *rap*

*+/+; UAS-fzrII.I/UAS-fzrII.I;+/+ x hs.FLP/Y; +/+; Act>CD2>GAL4 UAS-GFP/TM6b*

*+/+; UAS-fzrII.I/UAS-fzrII.I;+/+ x +/Y; +/+; Repo-GAL4/Tb*

#### C. *rap* alleles and Loco^p452^

*rap^E2^/rap^E2^; +/+; +/+ x +/+; +/+; Loco^P452^/Tb*

*rap ^E4^/rap^E4^; +/+; +/+ x +/+; +/+; Loco^P452^/Tb*

*rap ^E6^/rap^E6^; +/+; +/+ x +/+; +/+; Loco^P452^/Tb*

*rap ^x-2^/rap^x-2^; +/+; +/+ x +/+; +/+; Loco^P452^/Tb*

*rap ^x-3^/rap^x-3^; +/+; +/+ x +/+; +/+; Loco^P452^/Tb*

*w,rap^3^/w,rap^3^; +/+; +/+ x +/+; +/+; Loco^P452^/Tb*

*CS/CS; +/+; +/+ x +/+; +/+; Loco^P452^/Tb*

*w^1118^/w^1118^; +/+; +/+ x +/+; +/+; Loco^P452^/Tb*

*+/+; +/+; Loco^P452^/Tb*

*x +/+; +/+; Loco^P452^/Tb*

#### D. Loss of function *rap* alleles

*(111943) w[67c23] P{w[+mC]=lacW}rap[G0418] P{neoFRT} 19A/FM7c; P{ey-FLP.D}5/P{ey-FLP.D}5; +/+ x FRT19, Ubi-GFP, eyFLP1/Y; +/+; +/+*

*(111943) w[67c23] P{w[+mC]=lacW}rap[G0418] P{neoFRT} 19A/FM7c; P{ey-FLP.D}5/P{ey-FLP.D}5; +/+ x FRT19, UbiGFP/Y; +/+; hs.FLP/TM6B*

### Immunohistochemistry

Third instar larval eye discs and brains were dissected in PBS and fixed in 4% paraformaldehyde pH of 7.4. Brains were washed in PBS and incubated in primary antibodies for 2 hours at room temperature or overnight at 4°C. Primary antibodies used: Mab24B10 (Zipursky, Venkatesh, and Benzer, 1985) were used to visualize photoreceptor axons (1:5) supernatant, Developmental Studies Hybridoma Bank; Mab24B10 specifically stains photoreceptor cells in the retina and their axonal projections to the optic ganglion (Zipursky et al., 1985); anti-phospho-histone3 (Ser10); Phosphorylation on Ser 10 of histone H3 is a mitotic marker; anti-rabbit Mitosis Marker (Millipore, 1:100); ELAV-9F8A9-S anti-mouse, (Developmental Studies Hybridoma Bank, 1:5); anti-β-galactosidase polyclonal rabbit, Bioscience, (1:500), rabbit anti-GFP (Chemicon, 1:500); polyclonal rabbit (Chemicon, 1:500); rabbit anti-Repo polyclonal (Chemicon, 1:500). The tissues were washed in 1XPBS three times for 20 minutes and incubated in the secondary antibody for 90 minutes at room temperature. Secondary antibodies used: Fluorescein (FITC)-conjugated AffinityPure goat Anti-Mouse IgG (H+L) (1:100); Rhodamine (TRITC)-conjugated affinity pure goat anti-mouse (1:100); Rhodamine (TRITC)-conjugated affinity pure goat anti-rabbit (1:100); Cy5-conjugated affinity pure F(ab’)2 goat anti-rabbit (1:100); goat-anti rabbit FITC IgG (H+L) (1:100) Jackson-ImmunoResearch Laboratories. The tissues were washed three times with 1XPBT and mounted using Vectashield Mounting Medium, Vector Laboratories.

### Imaging and axon analysiss

The fluorescent samples were visualized using confocal microscope LSM 510 (Zeiss). LSM Image Browser was used to capture, filter and visualize z-stacks of the preparations. All images were visualized at 40x magnification 0.7 to 1.5 zoom and z-stack projections planes were compiled using LSM Image Browser. Each projection is a stack image composed of 20-30 focal planes, each advancing 1.5 μm along the z-axis. Images were edited with Adobe Photoshop CS3 and CS5. The lengths were quantified using projected images obtained from z-stack images generated using LSM Image Browser.

### Mosaic analysis

A mosaic analysis was performed by heat shocking first instar larvae for 15 or 30 min at 37°C water bath 24 or 48 hours after egg laying for gain of function experiments. The immunohistochemistry experiments were performed as described above. Images were edited with Adobe Photoshop CS3 and CS5.

### Statistical analysis

Analyses were performed with Prism 8 software (GraphPad Software). All statistical tests, significance was set to p < 0.05. The two-tailed unpaired Student’s t-test were used to analyze photoreceptor axon lengths or one-way ANOVA, where appropriate. Sample size (*n*) represents individual animals. All data are expressed at mean ± SEM.

## Results

### *rap* mutants exhibit axonal targeting abnormalities

The partial loss-of-function mutations in the *Drosophila rap* gene cause a rough external phenotype when compared to the wild type *Canton S*. It was previously shown that these mutations cause additional aberrant mitotic cell cycles in the developing eye (Pimentel and Venkatesh, 2005). Additional alleles of *rap; rap^x-2^; rap^x-3^; w, rap^3^*, not previously described, were examined for such defects and all showed aberrant mitotic cell cycle in the developing eye **(Figure 1)**. The *rap* mutants show an abnormal phospho-histone3 (Ser10) staining with cell division not restricted to the MF and differentiated regions of the eye disc **(Figure 1B-E)**. Examination of the protein expression patterns of Rap/Fzr showed that it is expressed in the nucleus of the post mitotic photoreceptor neurons; thus, it led us to investigate whether Rap/Fzr has a neuronal function (Pimentel and Venkatesh, 2005). The partial loss-of-function alleles in the *rap* mutants cause a range of severity in the external rough eye phenotype as compared to the laboratory wild type, *Canton S*. flies.

**Figure 1.**
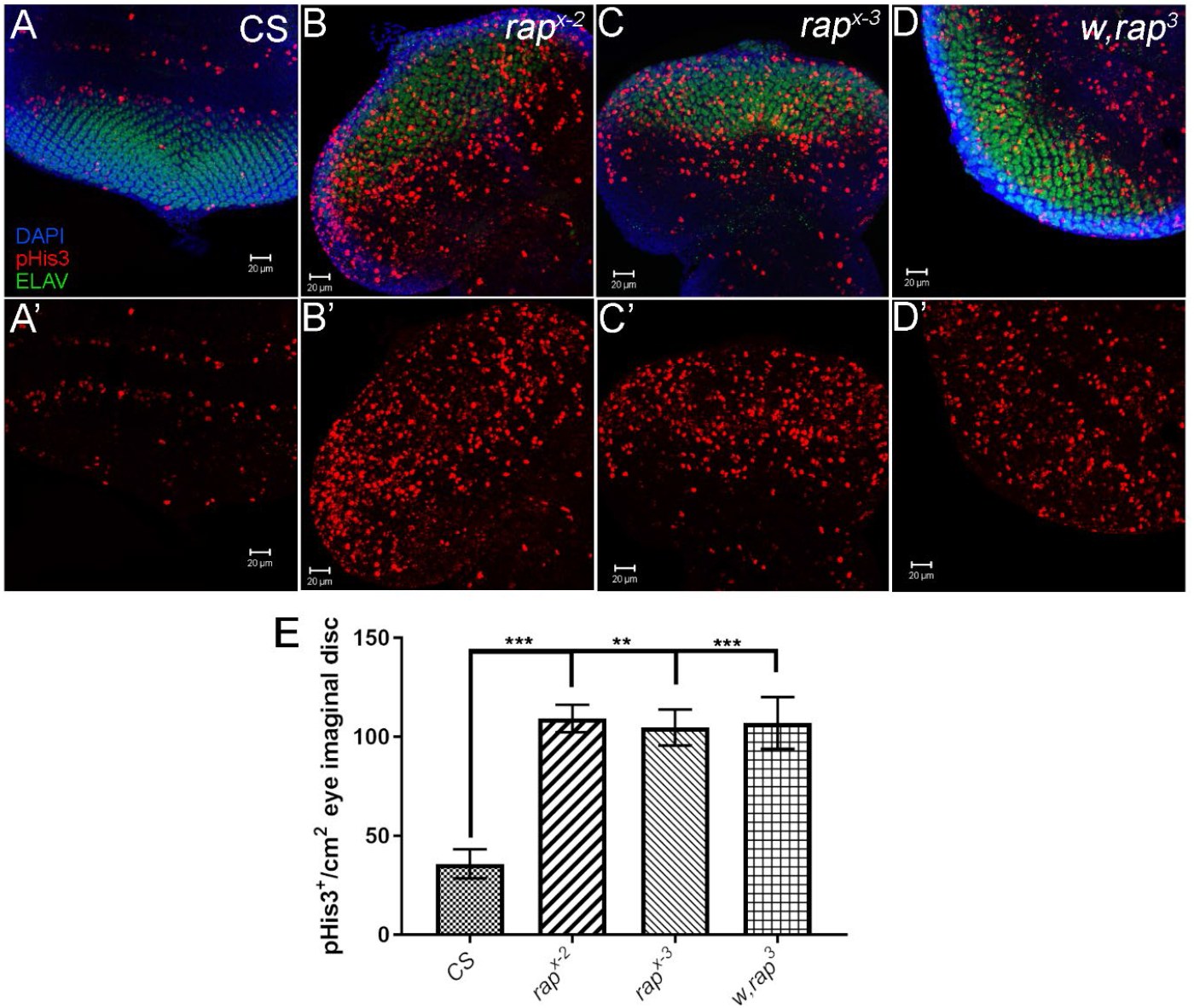
*rap* alleles stained for phospho-histone 3 (Ser10) show dysregulated mitotic cell division. Genotypes: A-A’) *CS/Y*; B-B’) *rap^x-2^/Y*, ***p = 0.0007; C-C’) *rap^x-3^/Y*, **p = 0.0011; D-D’) *w,rap^3^/Y*, ***p = 0.0009; E) Quantification of pHIS3^+^ cells/cm^2^ of eye imaginal disc, One-way ANOVA, alpha>0.05; [F(3, 9) = 0.2197, p = 0.8803]. Data represented as mean ± SEM; n = 3-4 animals per group. pHis3, red; ELAV, green; DAPI, blue. Scale bar 20 μm.

Thus, to test whether Rap/Fzr plays a role in the developing R cell neurons, we used development of *Drosophila* eye imaginal disk to examine axon projection patterns in developing eyes from *rap* partial loss-of-function mutants. Six *rap* alleles previously generated by X-rays (*rap^x-2^* and *rap^x-3^*), EMS (*rap^E2^, rap^E4^*, and *rap^E6^*), and P-element (*w,rap^3^*) mutagenesis by Tadmiri Venkatesh laboratory were used. The photoreceptor axons of third instar eye imaginal discs were analyzed for abnormalities in the axonal projection patterns using Mab 24B10, an antibody specific to photoreceptor cells and their axons in the eye imaginal disc (Zipursky et al., 1985). The wild type third instar eye discs of *CS and w^1118^* (*white* mutation, no pigment expression, previously shown to have normal photoreceptor axon projection patterns) flies show stereotypical axon projections from the eye imaginal disc, through the optic stalk, with properly organized lamina and medulla axons and clearly defined lamina plexus **(Figure 2A-B)**. Rap mutant alleles showed aberrant retinal axon projections. The different alleles of *rap* mutants examined showed allele specific variation in phenotype, with *rap^x-3^, rap^E4^*, and *rap^E2^*, **(Figure 2D-F)** showing photoreceptor axon overgrowth. All *rap* mutants show gaps in the lamina plexus, axonal clumping, disorganized lamina and medulla layers **(Figure 2C-H)**. The *rap* alleles show specific variations in the abnormal phenotype suggesting that Rap might operate in a gene dosage dependent manner. The alleles, *rap^x-2^, rap^x-3^*, and *w, rap^3^* **(Figure 2C-D, and H)** showed additional gaps in the lamina plexus, axon bundling in the medulla and lamina as well as the increase in the overall length of the projections **(Figure 2F-H)**.

**Figure 2.**
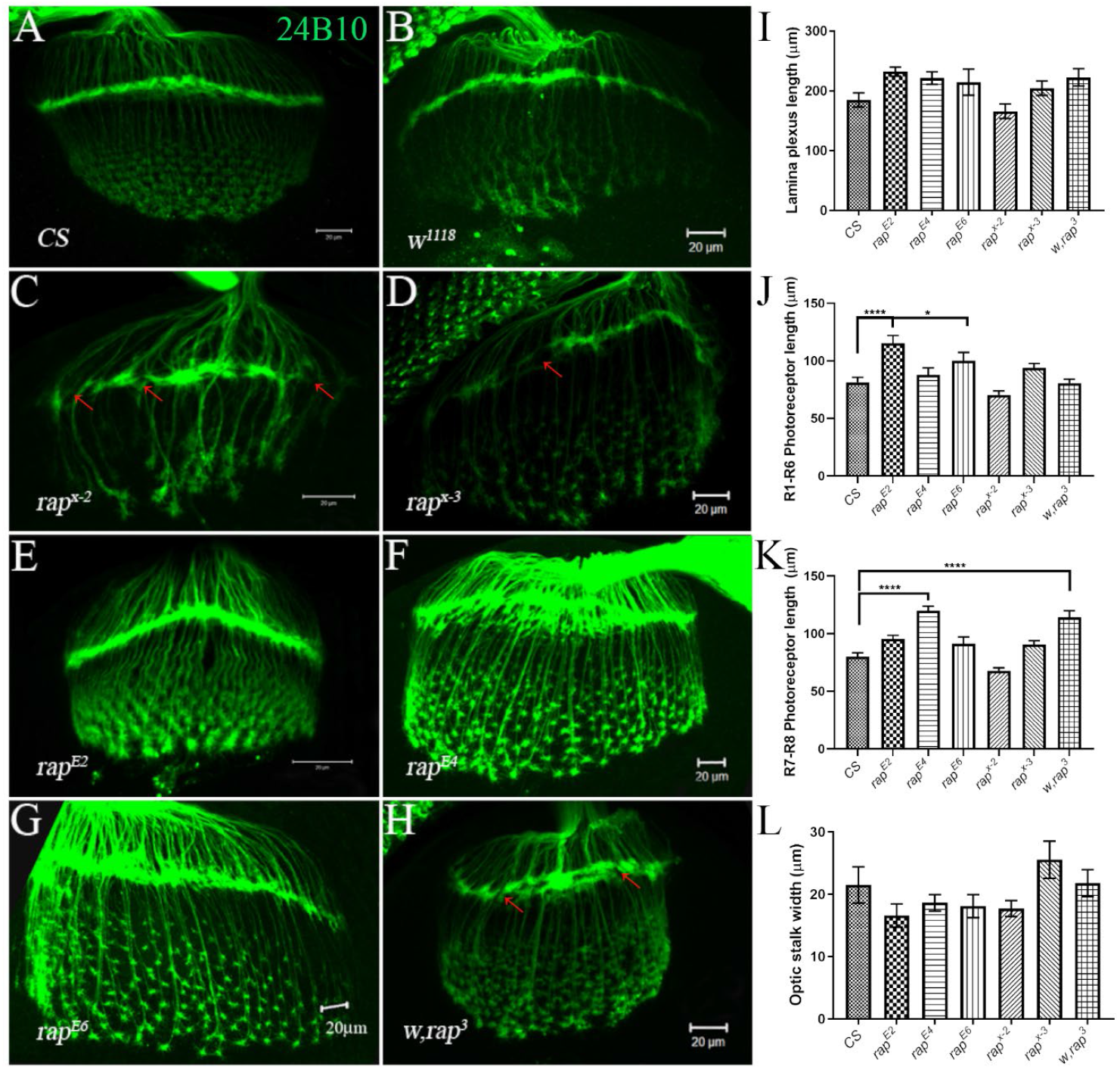
*rap/fzr* is required for proper development and target projection of photoreceptor axons. A-B) The wild type (*CS* and *w^1118^*) *Drosophila* eye shows proper innervations in the lamina and medulla. The axon projections do not show clumping or disorganization. They follow stereotypical patterns of targeting. C-H) *rap* mutants show allele specific abnormalities, lamina gaps (shown with arrows), disrupted medulla and axonal overgrowth. Axons stained with 24B10 in green. Quantification of photoreceptors of third instar larvae: I) Lamina plexus length (μm). One-way ANOVA, α > 0.05; [F(6, 58) = 2.486, p = 0.0329]; J) R1-R6 Photoreceptor length (μm). One-way ANOVA, α > 0.05; [F(6, 317) = 0.1359, p < 0.0001]. *p = 0.489, ****p < 0.0001; K) R7-R8 Photoreceptor length (μm). One-way ANOVA, α > 0.05; [F(6, 318 = 14.24, p < 0.0001]. *p = 0.0295. ****p < 0.0001; L) Optic stalk width (μm). One-way ANOVA, α > 0.05; [F(6, 58) = 1.678, p = 0.1428]. Data represented as mean ± SEM; n = 7-15 animals per group. Scale bar 20 μm.

Quantitative analysis of the *rap* allele defects in the photoreceptor axons also confirmed the phenotypic abnormalities. The results showed significant changes in the lengths R cell axons, *rap^E2^* **(Figure 2E, J)** and *rap^E6^* **(Figure 2G, J)** displaying a significantly longer R1-R6 R cells. While, *rap^E4^* **(Figure 2F, J)** and *w,rap^3^* **(Figure 2H, J)** showed a significantly longer R7-R8 R cell axons. Although the lengths of the lamina plexus are trending upwards for nearly all alleles, the differences were not statistically significant. There was no difference in the length of the optic stalk width, measured at the base of the disc, although some appear to be trending downwards. Additional parameters are needed to assess the severity of axonal projection patterns but due to clumping in the mutant flies the quantification of the number of photoreceptor axons in individual clumps could not be easily determined.

Our results are consistent with function of Rap/Fzr as an inhibitor of axon growth and elongation. The data is supported by the mammalian studies where the mammalian homolog Cdh1, was knocked down using siRNA in the axons in the cerebellar cortex showing growth and elongation (Konishi et al., 2004) and in rat cerebellar cells (Huynh et al., 2009). These results indicate that Rap/Fzr plays a role in the determination of axonal growth and inhibition of the axonal growth; thus, partial loss-of-function in the Cdh1 showed an increased length of the photoreceptor axons **(Figure 2)**.

### Photoreceptor cell axons R2-R5 show mistargeting in *rap* mutants

In order to determine which photoreceptor axon group causes the abnormal phenotype, we selectively labelled the R cell axons R2-R5 using a *ro-tau*-LacZ reporter (Fan et al., 2005) as the 24B10 antibody used to visualize *rap* mutants labels all R cells. The *ro-tau*-LacZ marker was crossed into the *rap* mutant background and the axon projections were examined. The wild type, *CS*, and the *ro-tau*-LacZ (parental strain) show well-defined lamina axons, stained with anti-β-galactosidase antibody **(Figure 3A)**. *rap* alleles show R2-R6 photoreceptor axons crossing the lamina plexus and terminating in the medulla. Also, in many instances R cells tend to clump at the lamina plexus, causing gaps **(Figure 3B-G)**. It was also noticed that *ro-tau*-LacZ reported was somewhat leaky and some of the expression was also seen in the R7-R8 cells, but the expression was at a lower level in the parental strain than in the double mutant cells. The data suggests that in *rap* partial loss-of-function alleles of *rap* cause developmental axonal changes, possibly changing the photoreceptor outgrowth or affecting proper R cell axon targeting.

**Figure 3.**
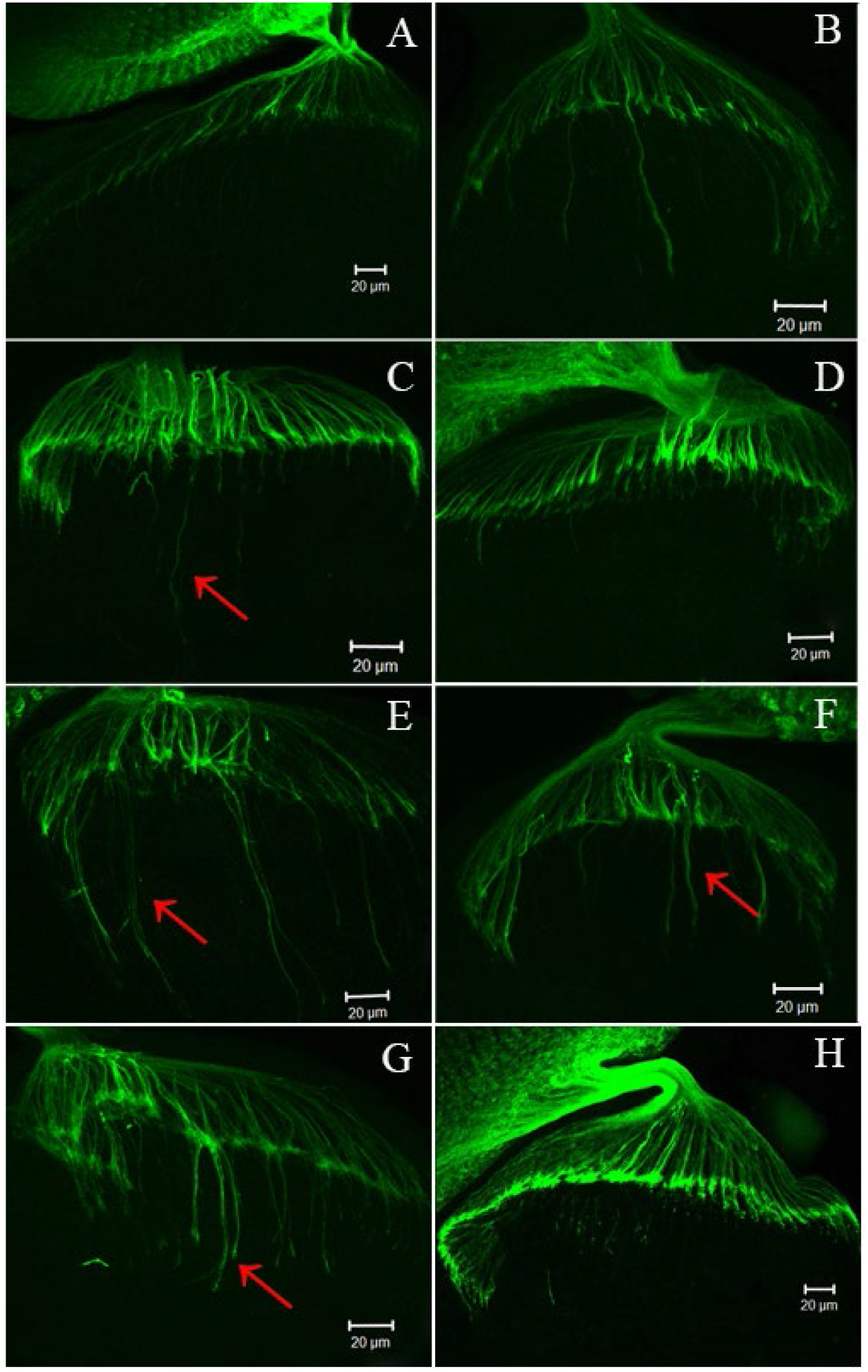
*rap/fzr* is required for proper R2-R5 photoreceptor targeting to the lamina in third instar optic lobe. A) *ro-tau-lacZ* parental strain; B) *rap^x-3^/Y; ro-tau-lacZ*; C) *rap^E2^/Y; ro-tau-lacZ*; D) *w,rap3^3^/Y; ro-tau-lacZ*; E) *rap^x-2^/Y; ro-tau-lacZ*; F) *rap^x-3^/Y; ro-tau-lacZ*; G) *rap^E4^/Y; ro-tau-lacZ*; H) *Loco^P452^/+; ro-tau-lacZ/+*. Red arrows point the R2-R5 cells bypassing the lamina plexus. 24B10, green. Scale bar 20 μm.

### Rap/Fzr functions cell autonomously in retinal axon targeting

*rap* is an essential gene and its expression is necessary for the survival of *Drosophila*, as null alleles are lethal. To assess whether Rap/Fzr functions in cell autonomous manner and to compare in vivo the phenotype of wild type and *rap* mutant in the retinal photoreceptor axons side by side, we conducted a mosaic analysis in a tissue- and time-specific manner using eye imaginal discs of third instar larvae. Eye-antenna discs are commonly used for the clone generation in a developing visual system (Xu and Rubin, 1993) and to assess axon targeting defects as reviewed elsewhere (Takeichi, 2007). Clone generation allows us to obtain homozygous null *rap-/rap*-mutant clones in the overall homozygous wild type, *rap+/rap+* twin spot, background.

To generate mosaic clones, we utilized a well described FLP-FRT system which allows for an efficient mitotic clone generation using the heat shock promoter driving Flippase expression acting on the FRT sites. FLP is a site-specific recombinase which recognizes FRT (FLP recombination target) sites (Theodosiou and Xu, 1998). This site-specific recombination system allows for a deletion of a particular DNA fragment located between FRT sites, in this case to remove *rap* allele. *Heat shock* and *eye-Flp* promoters were used to express the Flippase enzyme. GFP fusion reporter was used to localize and visualize the clones.

Our data indicates that the regions of the *rap-/rap*-clones, identified by the absence of GFP staining, show aberrant axon projections **(Figure 4A)** and axonal clumping in the eye disc **(Figure 4B)**. Clumping of axons and various gaps in the lamina plexus were observed in the clone patches but the general morphology of the surrounding cells was normal. The phenotype appears cell autonomous since the surrounding genotypically heterozygous or wild type tissue was phenotypically normal.

**Figure 4.**
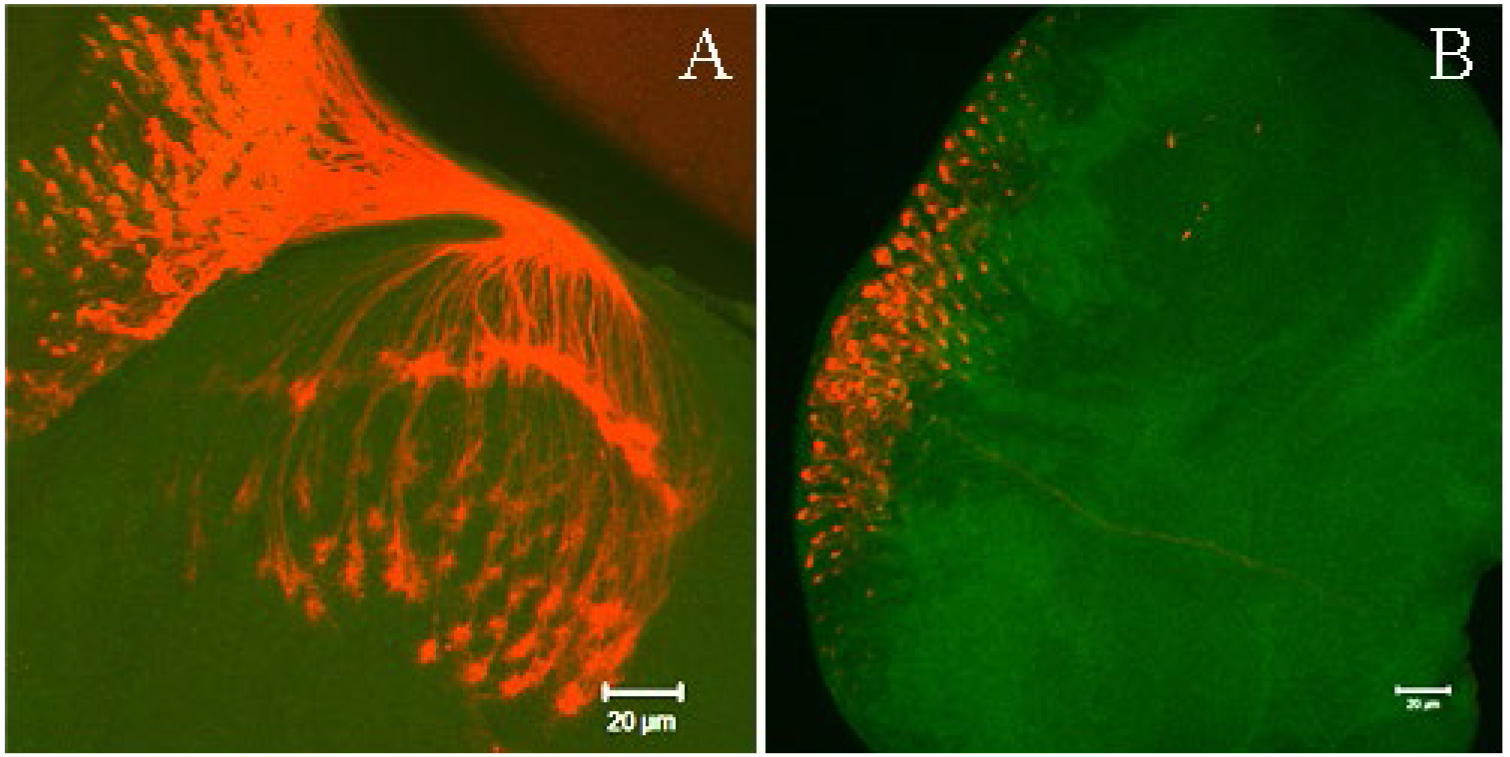
Loss of function clones of *rap-/rap*-using FRT site specific recombination show *rap* photoreceptor defects. Genotype: *w67c23 P{lacW}rapG0418 P{neoFRT}19A/FRT19,UbiGFP,eyFLP; P{ey-FLP.D}5/+*. The outlines point to the areas where *rap-/rap-* clones were generated in the eye disc, showing abnormal photoreceptors. Arrow points to the gaps in the lamina plexus. 24B10, red; GFP, green. Scale bar 20 μm.

To confirm the effects seen in ey-FLP animals we generated additional clones using two different reporter lines **(Figure 5A-D, I-L)** with Hsp-FLP located on different chromosomes to test the signal strength and consistency. The clones are oputlined with a white line **(Figure 5E-I, M-P)**. The visualization of clones in the third instar larva poses a challenge, as only the clones in the optic lobe surface can be seen; thus, the images do not show clones on the interior of the lobe. This explains why axons in other brain regions besides the enclosed clone areas **(Figure 5)** have abnormal axonal patterns, forming clumps, gaps and disorganized lamina plexus. These data suggest that Rap/Fzr functions in cell autonomous manner. Additional experiments need to be performed for a more definitive conclusions such as MARCM, mosaic analysis with a repressible marker, to allow visualization of a single cell clone in the heterozygous wild type background (Lee and Luo, 2001).

**Figure 5.**
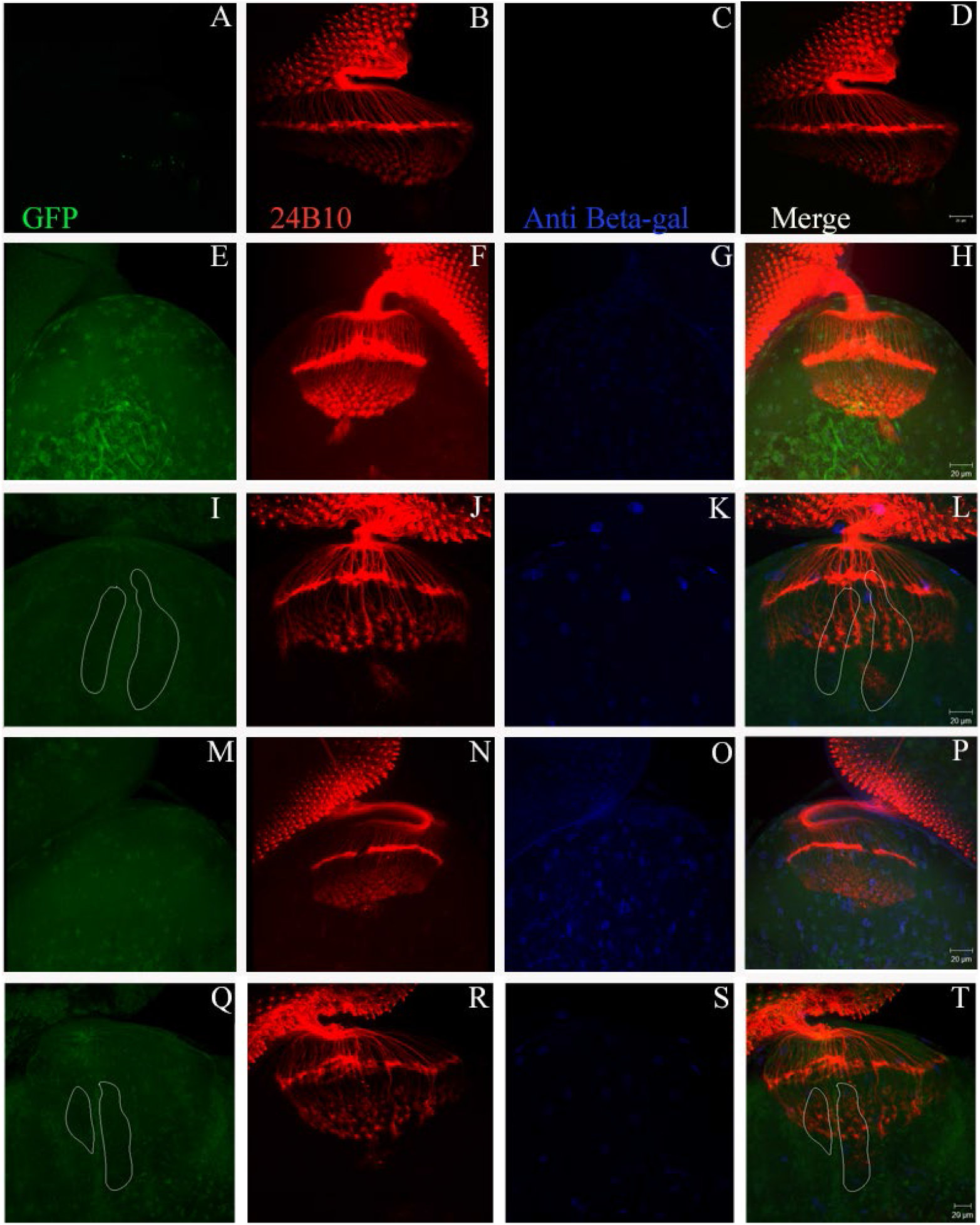
The loss-of-function *rap* clones with abnormal photoreceptors. A-D and I-L. Show parental strains used to generate the clones. The image shows axonal clumping, mistargeting of photoreceptor axons, disorganized lamina plexus and the overall morphology of the clones is affected. Images D, H, L, and P are merges of GFP, 24B10, and anti-β-galactosidase staining. The control strain genotype are: A-D) *FRT19,UbiGFP,eyFLP1/Y;+/+; +/+;* and I-L) *FRT19,UbiGFP/Y; +/+; hs.FLP/+;* While the mutant closed are represented in E-H. and M-P. Loss-of-function clones outlined in white. Genotypes: E-H) *w67c23 P{lacW}rapG0418 P{neoFRT}19A/FRT19,UbiGFP,hs.FLP1; P{ey-FLP.D}5/+; +/+;* and M-P) *w67c23 P{lacW}rapG0418 P{neoFRT}19A/FRT19,UbiGFP; P{ey-FLP.D}5/ +; hs.FLP/+*. GFP, green; 24B10, red; anti-β-Gal, blue. Scale bar 20 μm.

### Gain-of Function experiments show that Rap/Fzr is required for proper axon targeting

Since *rap* is required for proper R cell axonal length regulation as shown in the loss-of-function experiments, we wanted to examine whether the gain-of-function would cause the opposite effect, axon shortening. Thus, we asked whether the gain-of-function in Rap/Fzr modulates proper axon targeting and length. To test the effects of overexpression of Rap/Fzr, gain-of-function *rap/fzr* clones were generated and screened for photoreceptor axon projection using: UAS/GAL4 (Duffy, 2002) combined with FLP/FRT (Golic et al., 1997). GAL4 is a transcriptional activator from yeasts (McGuire et al., 2004), and is expressed in tissue specific manner while UAS is a transgene, an upstream activator sequence (St Johnston, 2002). The gain-of-function study would further ensure that the protein in question is playing a role in the alteration of the phenotype. While, UAS/GAL4 system allows direct assessment of the effect of expression of a gene of interest with the ability to report and visualize its activity.

The *rap* gene was cloned downstream of a UAS sequence and the gene was expressed in the presence of GAL4 protein that binds to the UAS to activate transcription. Both GAL4 and *UAS-rap/fzr* were carried in different strains, allowing the parental viability. The experiment utilized the *UAS-fzrII.I, UAS-GFP* as a visible reporter gene, and combines the UAS/GAL4 system with heat shock FLP and Actin>CD2>FRT-GAL4; UAS-GFP cassette, (where > are FRT sequences). Under a heat shock, Flippase will be activated causing site-specific recombination of the FRT sequences and excision of the FRT>CD2 flanking sequence. The excision will allow the expression of GAL4 from the Actin promoter and binding of the GAL4 protein to the UAS sequences and activating the expression of *rap/fzr* and GFP reporter gene. All cells labeled with GFP will also overexpress Rap/Fzr protein allowing for the visualization of the affected cells. The heat shock was administered 24 **(Figure 6A-D)** or 48 hours **(Figure 6E-P)** after egg laying at 37°C water bath. The gain of function clones show axon clumping and axon shortening in the early induced clones **(Figure 6A-D)** as well as in the later induced clones **(Figure 6E-P)**.

**Figure 6.**
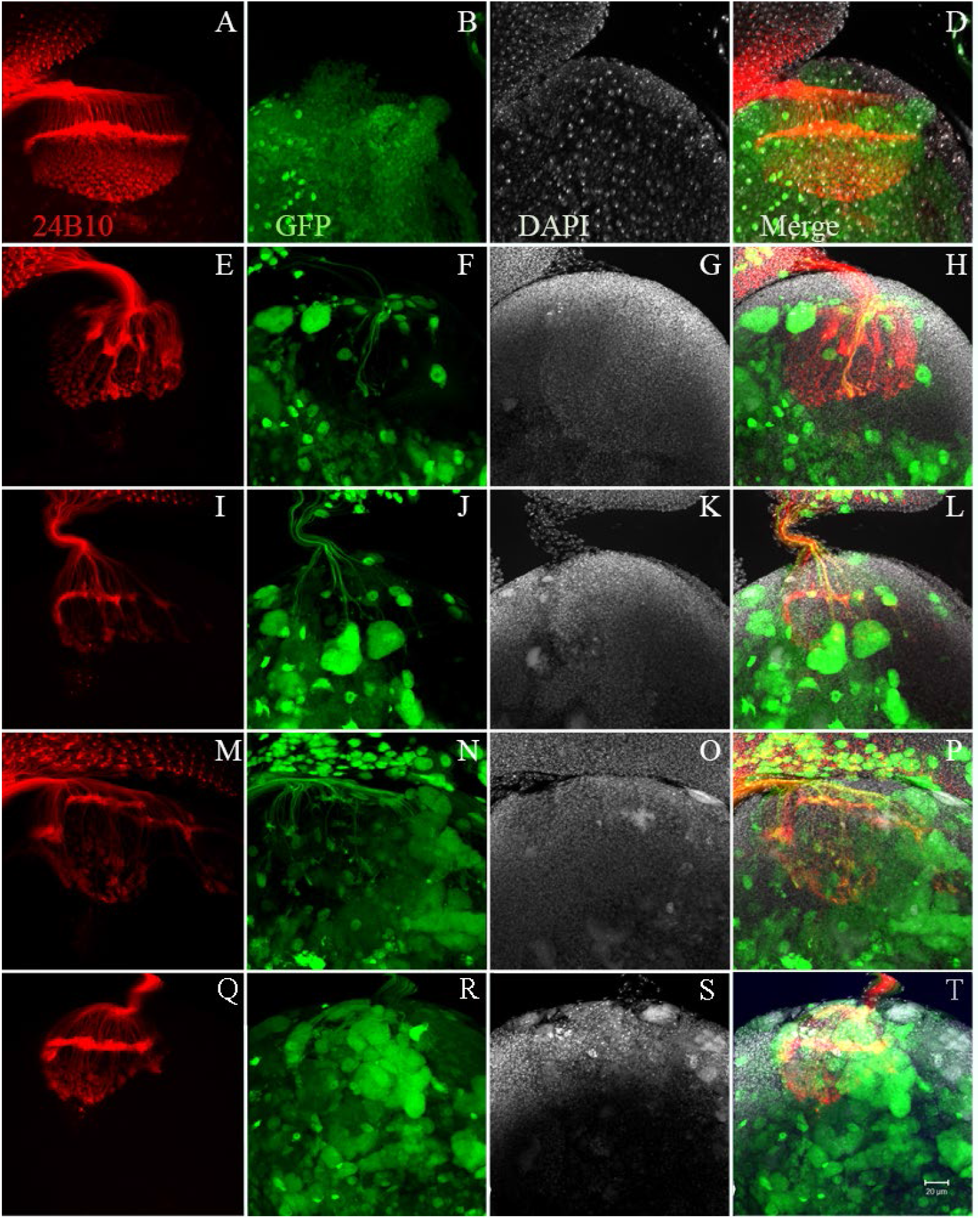
Gain-of-function *rap* clones expressing GFP and FZR show R cell axon defects. The gain of function clones show axon clumping, axonal bundles and shortening of axons in early induced clones. A-D) heat shocked 24 hours after egg laying; E-P) heat shocked 48 hours after egg laying. D) is a merge of A-C; H) merge of E-G; L) merge of I-K; and P) merge of L-O. Genotype: *hsp.FLP/+; UAS-fzr II.I/+; Act>ΔGa14 UAS-GFP/+*. 24B10, red; GFP, green ΔDAPI, white. Scale bar 20 μm.

Thus, the gain-of-function of *rap/fzr* in the eye imaginal disc causes axonal clumping, disorganization of lamina plexus displayed many gaps. These data suggest that Rap/Fzr is necessary for proper axonal targeting and proper development of the *Drosophila* visual system in the third instar larvae. Interestingly, the lamina plexus shows clusters of R cells fluctuating from the proper orientation in a step-wise fashion. It was previously shown that gain-of-function of *rap* causes an increase in the neuroblast and decrease in the glia numbers in the optic lobe of *Drosophila* (Kaplow et al., 2008). It suggested that the overexpression of Rap causes an increased number of neurons that cannot fit onto the plane of the lamina plexus and cause the disorganization of that region. Although, the photoreceptor axons were not measured, the length of the R cells is significantly decreased.

### Loco dominantly interacts with *rap* to regulate axon targeting

Previous studies have shown that Rap, a ubiquitin ligase, targets Loco for ubiquitination and degradation (Kaplow et al., 2008). Loco was previously shown to affect the number of neurons and glia (Kaplow et al., 2008); thus, we hypothesized that it may play a role in axonal targeting. Thus, we decided to investigate possible interactions of Rap and Loco in axonal targeting by asking whether Loco dominantly interact with *rap* to regulate axon targeting.

The *loco^P452^* allele is a homozygous lethal with rare third instar escapers that are characterized by locomotion problems and pupal lethality (Yu et al., 2005). Third instar larvae escapers were stained for photoreceptor axons **(Figure 7B)** and showed overgrowth of R1-R6, R7-R8, as well as an elongated lamina plexus **(Figure 7I-L)** but no difference in the optic stalk width **(Figure 7K)**. The cross to *ro-tau*-LacZ reported did not show mistargeting or R2-R6 into the medulla (not shown), thus Loco may act in a different pathway in the normal *rap* background.

**Figure 7.**
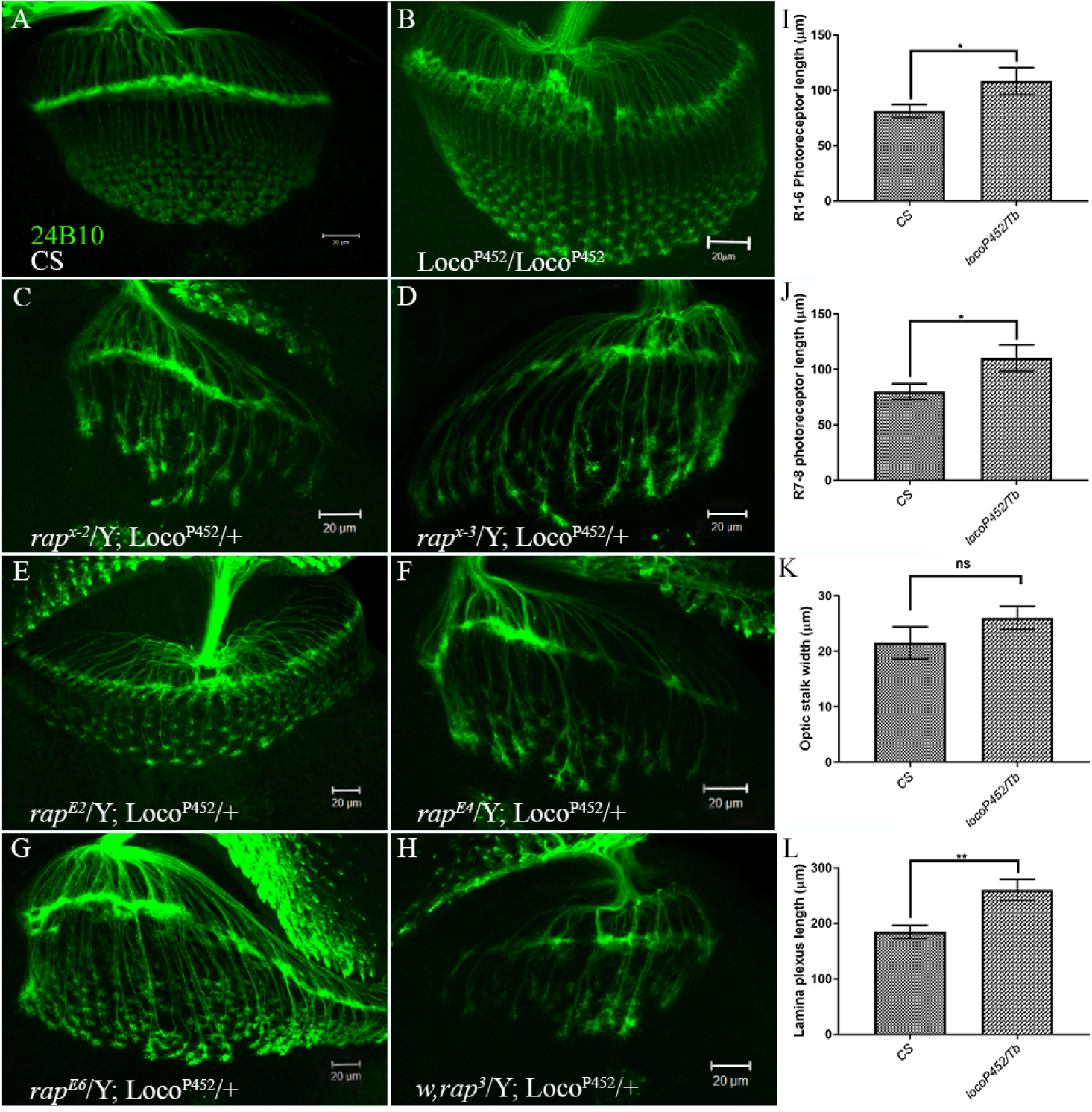
*Loco^P452^* enhances photoreceptor axon defects in *rap* allele background. The *rap* phenotype is enhanced by the single copy of Loco^P452^. Axons show an increased bundling, more lamina plexus gaps and overall abnormal morphology of axonal projections. 24B10, green. A) CS; B) *Loco^P452^/Loco^P452^; C) rap^x-2^/Y; Loco^P452^/+; D) rap^x-3^/Y; Loco^P452^/+; E) rap^E2^/Y; Loco^P452^/+; F) rap^E4^/Y; Loco^P452^/+; G) rap^E6^/Y; Loco^P452^/+; H) w,rap^3^/Y; Loco^P452^/+. I-J)* Bar graphs representing Loco^P452^/Tb allele R cell photoreceptor measurements. The measurements show the lengths of the following: I) R1-6 photoreceptor length, *p = 0.0480; J) R7-8 photoreceptor length, *p = 0.0385; K) optic stalk, p = 0.2441; L) Lamina plexus length, **p = 0.0026. Two-tailed unpaired student t-test, alpha<0.05. Data represented as mean ± SEM; n = 8-10 animals per group. Scale bar 20 μm.

To study a possible interaction of *rap/fzr* with the *loco^P452^* allele, we crossed different *rap* alleles to loss-of-function allele *loco^P452^* to generate double mutants that contained single copy of *loco^P452^*. Third instar larvae eye imaginal discs were examined for the R cell axonal projection defects. Surprisingly, *loco* which was previously shown to dominantly suppress rough eye phenotype in the adult *Drosophila* eye in genetic modifier screen (Kaplow et al., 2007), enhanced the axon bundling and mistargeting of the R axons in the lamina, medulla and lamina plexus in the third instar larvae when crossed to *rap* mutants. The lamina plexus shows an increased number of gaps and axon bundling than the single mutant *rap* alleles **(Figure 7C-H)**.

These results suggest that *rap/fzr* plays a role in the axonal development, targeting and length, while *loco* helps to coordinate the axonal length. It also suggests that Loco and Rap may be involved in another pathway that allows for a cross-talk since double mutant Rap/Loco do not show the same phenotype but an enhancement of the *rap* phenotype.

### Liprin-α interacts with Rap/Fzr to regulate axon targeting

Previous studies have shown that a scaffolding protein Liprin-α is required cell autonomously in all R cells for proper retinal axon target selection (Choe et al., 2006b; Prakash et al., 2009). Thus, we wanted to determine whether Liprin-*α* genetically interacts with Rap/Fzr to enhance the R cell axonal defects in third instar *Drosophila* eye imaginal disc. To test for this interaction, we crossed *rap* allele mutants to two mutants of *Liprin-α* generated by P-element mutagenesis that disrupt the open reading frame of the *Liprin-α* gene. The results have shown that both of the heterozygous deficiencies did not exhibit any gross axonal defect phenotypes **(Figure 8A,H)**. When *Liprin-α* loss-of-function alleles were crossed to 6 different alleles of *rap*, in nearly all cases there was an enhancement of axonal clumping, mistargeting that was allele specific **(Figure 8B-G, I-N)**. Interestingly, double mutants that showed significant difference in R1-6 or R7-8 in the individual *rap* mutants had the most severe defects **(Figure 8C, D, G, J, K, N)**. Interestingly, the less severe single *rap* mutants, rap^x-2^ and rap^x-3^, showed a severe enhancement of the axon targeting defects, with increased axon clumping, lamina plexus gaps, and improper axon extensions **(Figure 8F, L-M)**. Similarly, the alleles of *rap, rap^E2^, rap^E4^, rap^E6^, and w,rap^3^*, that displayed a more severe phenotypes, had an enhancement of the axonal defects, with an exception for the milder allele of *rap^E2^* **(Figure 8B-D, G, I-K, N)**.

**Figure 8.**
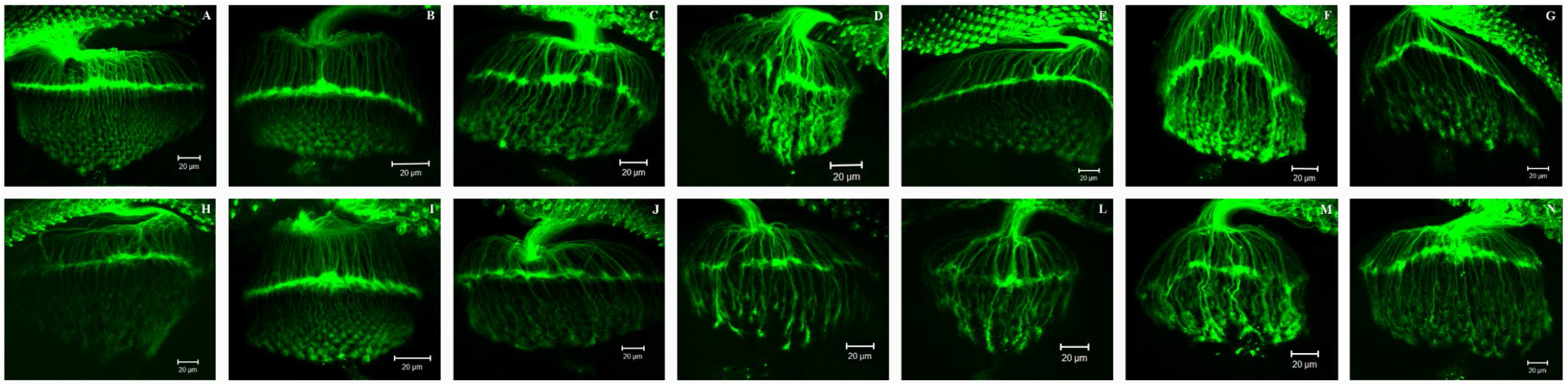
Liprin-α deficiencies enhance photoceptor axon defects in *rap* allele background. A) *+/Y; P{y[+t7.7] w[+mC]=wHy}Liprin-alpha[DG23609]/+; +/+* B) *rap^E2^/Y; P{y[+t7.7] w[+mC]=wHy}Liprin-alpha[DG23609]/+; +/+* C) *rap^E4^/Y; P{y[+t7.7] w[+mC]=wHy}Liprin-alpha[DG23609]/+; +/+* D) *rap^E6^/Y; P{y[+t7.7] w[+mC]=wHy}Liprin-alpha[DG23609]/+; +/+* E) *rap^x-2^/Y; P{y[+t7.7] w[+mC]=wHy}Liprin-alpha[DG23609]/+; +/+* F) *rap^x-3^/Y; P{y[+t7.7] w[+mC]=wHy}Liprin-alpha[DG23609]/+; +/+* G) *w,rap^3^/Y; P{y[+t7.7] w[+mC]=wHy}Liprin-alpha[DG23609]/+; +/+* H) *+/Y; P{w[+mC] y[+mDint2]=EPgy2}Liprin-alpha[EY21217]/; +/+* I) *rap^E2^/Y; P{w[+mC] y[+mDint2]=EPgy2}Liprin-alpha[EY21217]/; +/+* J) *rap^E4^/Y; P{w[+mC] y[+mDint2]=EPgy2}Liprin-alpha[EY21217]/; +/+* K) *rap^E6^/Y; P{w[+mC] y[+mDint2]=EPgy2}Liprin-alpha[EY21217]/; +/+* L) *rap^x-2^/Y; P{w[+mC] y[+mDint2]=EPgy2}Liprin-alpha[EY21217]/; +/+* M) *rap^x-3^/Y; P{w[+mC] y[+mDint2]=EPgy2}Liprin-alpha[EY21217]/; +/+* N) *w,rap^3^/Y; P{w[+mC] y[+mDint2]=EPgy2}Liprin-alpha[EY21217]/; +/+*

These results suggest that *rap/fzr* and *Liprin-α* act on the similar processes that are required for proper axon targeting in the developing fly eye. These results cannot confirm whether the two interact biochemically but further experiments would be necessary to assess that possibility.

## Discussion

The work presented in this paper was conducted to test whether Rap/Fzr, a *Drosophila* homolog of the mammalian Cdh1 plays a role in photoreceptor axon targeting in *Dropospila* third instar larvae eye imaginal disc. The results showed that *rap* plays a significant role in axon length determination as *rap* alleles showed abnormal axonal projections of R2-R5 as well as R7-R8, with clumping or photoreceptor axons, abnormal lamina, lamina plexus and medulla morphology, and axon overextension. *rap* alleles showed allele specific variations in the abnormal phenotype, suggesting that Rap operates in a gene dosage dependent manner and may affect multiple biochemical pathways regulating axon targeting. This data is consistent with previously described mammalian data (Konishi et al., 2004). Thus, *rap* normally functions to inhibit axonal growth.

Loss-of-function mosaic studies utilizing FLP/FRT system suggest that Rap/Fzr acts in a cell autonomous manner in retinal axon targeting, as only *rap-/rap-* clones showed aberrant axon projections, clumping of axons, and various gaps in the lamina plexus, but the general morphology of the surrounding R cells appeared normal. Additional experiments are required to determine which R cells are specifically affected. One possible way would be to use MARCM, mosaic analysis with a repressible marker, that allows visualization of a single cell clone in the heterozygous wild type background (Lee and Luo, 2001; Luo, 2007).

In addition, gain-of-function experiments utilizing GAL4/UAS and FLP/FRT systems showed that Rap/Fzr modulates proper axon targeting and length. Gain-of-function of *rap/fzr* in the eye imaginal disc caused axonal shortening, clumping, and disorganization of lamina plexus, showing clusters of R cells improperly orientation in a step-wise fashion. It suggests that the overexpression of Rap most likely leads to an increased number of neurons or glia that cannot fit onto the plane of the lamina plexus and cause the disorganization of that region. These data suggest that proper dosing of Rap/Fzr is required for proper axonal targeting and development of the *Drosophila* visual system.

Axonal development is regulated by multiple signal transduction cascades that require proper scaffolding proteins to mediate the necessary protein interactions. Thus, to test for additional Rap interactions, we generated double mutants by crossing our *rap* alleles with two scaffolding proteins, Loco or Liprin-α loss-of-function mutations. Previous studies have shown that Rap targets Loco for ubiquitination and degradation (Kaplow et al., 2008), thus we decided to investigate possible interactions of Rap and Loco in axonal targeting by asking whether Loco dominantly interact with *rap* to regulate axon targeting. Loco was previously shown to affect the number of neurons and glia (Kaplow et al., 2008); thus, it may play a role in axonal targeting. Surprisingly, *loco* which was previously shown to dominantly suppress rough eye phenotype in the adult *Drosophila* eye in genetic modifier screen (Kaplow et al., 2007), enhanced the axon bundling and mistargeting of the R axons in the lamina, medulla and lamina plexus in the third instar larvae. The lamina plexus showed an increased number of gaps and axon bundling than the single mutant *rap* alleles. These results suggest that interaction of Rap/FZR and Loco plays a significant role in the axonal development, targeting and length determination of R cells. It also suggests that Loco and Rap may be involved in another pathway that allows for a cross-talk, as double mutant Rap/Loco showed an enhancement of the axonal defects.

In addition, Liprin-α has been shown to be involved in photoreceptor axonal targeting (Choe et al., 2006a; Hofmeyer et al., 2006); yet, whether Rap/Fzr and Liprin-α interact in retinal axon targeting was unknown. Thus, we investigated whether Rap/Fzr interacts with Liprin-α in axon targeting in the third instar larvae. *rap/fzr*/Liprin-α double mutant crosses also showed an enhancement of axonal phenotype defects, axonal clumping, disorganized lamina and medulla with gaps in the lamina plexus. Our results show that the partial loss-of-function *rap* phenotype is enhanced by a single copy of the mutant Liprin-α. This is consistent with a role for Rap/Fzr in the degradation of Liprin-α as previously suggested (Choe et al., 2006a). Thus, it is possible that Rap and Liprin-α may be involved in a cross-talk regulating axon targeting.

Additional biochemical data is needed to further assess function of in *rap* and its interactions with Loco and with Liprin-α. Future studies should investigate the axonal lengths of the double mutant crosses with *rap* and additional Rap targets. Axonal elongation may play a role in treating axonal regeneration, thus additional experiments that would test the *rap* effect in the injury may shed a light on the axonal regeneration properties. Physiological and behavior studies are required to assess the functional aspects of these defects, and should assess their role in other neuronal subtypes.

## Conclusions

In conclusion, our study showed that *rap* is required in R cells for proper targeting of the developing axons to the appropriate layers of the lamina and medulla. We also showed that loss of *rap* in combination with the loss of *loco* or Liprin-α significantly enhances the axonal defects in the third instar developing fly eye, suggesting that all three genes are required for the proper development of R cell axonal projections.

## Authors’ contributions

Both authors contributed equally to this work.

## Acknowledgements

This work was supported by: The City College Fellowship and NIH.

